# Autophagy is required for the rejection of self-incompatible pollen in two accessions of transgenic *Arabidopsis thaliana*

**DOI:** 10.1101/2020.08.26.268326

**Authors:** Stuart R. Macgregor, Hyun Kyung Lee, Hayley Nelles, Daniel C. Johnson, Tong Zhang, Chaozhi Ma, Daphne R. Goring

**Affiliations:** Department of Cell & Systems Biology, University of Toronto, Toronto, Canada M5S 3B2; National Key Laboratory of Crop Genetic Improvement, National Center of Rapeseed Improvement in Wuhan, Huazhong Agricultural University, Wuhan 430070, China; Centre for the Analysis of Genome Evolution & Function, University of Toronto, Toronto, Canada M5S 3B2

**Keywords:** Autophagy, self-incompatibility, pollen-pistil interactions, reproduction, *SP11/SCR*, *SRK*, *ARC1*, *ATG7*, *ATG5*, *GFP-ATG8a*

## Abstract

Successful reproduction in the Brassicaceae is mediated by a complex series of interactions between the pollen and the pistil, and some species have an additional layer of regulation with the self-incompatibility trait. While the initial activation of the self-incompatibility pathway by the pollen S-locus protein11/*S*-locus cysteine-rich peptide and the stigma *S* Receptor Kinase is well characterized, the downstream mechanisms causing self-pollen rejection are still not fully understood. In previous studies, we had detected the presence of autophagic bodies with self-incompatible pollinations in *Arabidopsis lyrata* and transgenic *A. thaliana* lines, but it was not known if autophagy was essential for self-pollen rejection. Here, we investigated the requirement of autophagy in this response by crossing mutations in the essential *AUTOPHAGY7* (*ATG7*) or *AUTOPHAGY5* (*ATG5*) genes into two different transgenic self-incompatible *A. thaliana* lines in the Col-0 and C24 accessions. By using these previously characterized transgenic lines that express *A. lyrata* and *A. halleri* self-incompatibility genes, we demonstrated that disrupting autophagy can weaken their self-incompatible responses in the stigma. When the *atg7* or *atg5* mutations were present, an increased number of self-incompatible pollen were found to hydrate and form pollen tubes that successfully fertilized the self-incompatible pistils. Additionally, we confirmed the presence of GFP-ATG8a labelled autophagosomes in the stigmatic papillae following self-incompatible pollinations. Together, these findings support the requirement of autophagy in the self-incompatibility response and add to the growing understanding of the cellular events that take place in the stigma to reject self-pollen.

**One Sentence Summary:** In self-incompatible transgenic *Arabidopsis thaliana* lines, autophagy is an integral part of the cellular responses in the stigma to efficiently block fertilization by self-incompatible pollen.

## Introduction

Reproduction in the Brassicaceae is a tightly controlled process, with many components contributing to successful pollen-pistil interactions (reviewed in Doucet et al., 2016; Mizuta and Higashiyama, 2018; Johnson et al., 2019; Adhikari et al., 2020). The Brassicaceae dry stigma lacks surface secretions and instead regulates the delivery of resources to the pollen shortly after pollination. This leads to the selective hydration of compatible pollen grains, followed by the penetration of the pollen tube into the stigmatic barrier to grow down the reproductive tract (reviewed in Dickinson, 1995; Palanivelu and Tsukamoto, 2012; Doucet et al., 2016; Adhikari et al., 2020). It is also at this initial cell-cell communication stage that self-incompatible pollen is recognized and blocked by the stigma to prevent inbreeding (reviewed in Jany et al., 2019; Nasrallah, 2019; Durand et al., 2020). A recent live-cell imaging study in *Arabidopsis thaliana* compared pollen hydration rates of compatible and self-incompatible pollen grains and found that while compatible pollen grains rapidly hydrated, self-incompatible pollen failed to fully hydrate which then led to a failure in pollen germination or a failure of pollen tubes to penetrate the stigmatic papilla cell wall (Rozier et al., 2020). Furthermore, actin foci were present in the stigmatic papillae at the site of emerging pollen tubes for compatible pollen only. The actin focalization is likely associated with the stigmatic papilla cellular responses to enable compatible pollen hydration and pollen tube penetration (Rozier et al., 2020).

The Brassicaceae self-incompatible response is initiated by the binding of the *S*-haplotype-specific pollen ligand, *S*-locus protein 11/*S*-locus cysteine-rich peptide (SP11/SCR) to the corresponding stigma-specific *S* Receptor Kinase (SRK). This leads to the phosphorylation of the SRK kinase domain and activation of the downstream signaling responses, ultimately leading to the rejection of the self-incompatible pollen grain reviewed in Jany et al., 2019; Nasrallah, 2019). Not all Brassicaceae species possess this trait due to mutations in the key self-incompatibility genes, *SP11/SCR* and *SRK* (reviewed in Durand et al., 2020). For example, the transition of *A. thaliana* to a fully selfing species is associated with mutations in the *SCR* and *SRK* genes (Kusaba et al., 2001; Shimizu et al., 2008; Boggs et al., 2009; Tsuchimatsu et al., 2010; Guo et al., 2011; Tsuchimatsu et al., 2017). In contrast, other family members like *A. lyrata* and *A. halleri* have intact *SCR* and *SRK* genes, which result in a robust self-incompatibility response upon self-pollination (Kusaba et al., 2001; Schierup et al., 2001; Goubet et al., 2012).

A number of studies have reintroduced the self-incompatibility trait back into *A. thaliana* by introducing *A. lyrata* or *A. halleri SP11/SCR* and *SRK* transgenes (Nasrallah et al., 2004; Boggs et al., 2009; Indriolo et al., 2014; Iwano et al., 2015; Zhang et al., 2019; Rozier et al., 2020). However, the success of this approach was accession-dependent; for example, transgenic experiments in *A. thaliana* Col-0 failed to produce self-incompatible plants, whereas experiments using *A. thaliana* C24 were very successful (Nasrallah et al., 2002; Boggs et al., 2009; Iwano et al., 2015; Rozier et al., 2020). The cause of this was linked to inverted repeats within the *A. thaliana SRKA* pseudogene which triggers RNA silencing of the introduced *SRK* transgenes (Fujii et al., 2020). For the Col-0 accession, the inverted repeat was found in a conserved region of the predicted *pseudoSRKA* kinase domain resulting in the silencing of closely related *SRK* transgenes. For the C24 accession, the inverted repeat was found in a more polymorphic region of the predicted *pseudoSRKA* ectodomain and appeared to be less effective at inducing RNA silencing of related *SRK* transgenes (Fujii et al., 2020).

We have found that a strong self-incompatibility response can also be recapitulated in *A. thaliana* Col-0 when the *A. lyrata* or *A. halleri ARC1* transgene was added along with the *A. lyrata* or *A. halleri SP11/SCR* and *SRK* transgenes (Indriolo et al., 2014; Zhang et al., 2019). The ARC1 E3 ubiquitin ligase is proposed to act directly downstream of SRK in the self-incompatibility signaling pathway (Gu et al., 1998; Stone et al., 1999; Stone et al., 2003). A large part of the *ARC1* gene has been deleted in the *A. thaliana* genome, and knockdowns of *ARC1* in self-incompatible *Brassica napus* and *A. lyrata* resulted in a partial breakdown of self-incompatibility (Stone et al., 1999; Indriolo et al., 2012). ARC1 is proposed to target cellular components that are required in the stigma to accept compatible pollen, such as the EXO70A1 exocyst subunit (Samuel et al., 2009) and *Brassica* GLO1 (Sankaranarayanan et al., 2015). Self-incompatible pollinations also induced a strong and rapid cytosolic calcium influx in the stigmatic papillae (Iwano et al., 2015). These combined self-incompatibility responses are proposed to disrupt the secretory activity in the stigmatic papillae which would be required for compatible pollen acceptance (Dickinson, 1995; Elleman and Dickinson, 1996; Samuel et al., 2009; Safavian and Goring, 2013; Safavian et al., 2015; Goring, 2017).

In previous work, we uncovered signs of autophagy activation in *A. lyrata* and *A. thaliana* stigmatic papillae within 10 minutes of self-incompatible pollinations, but not with compatible pollinations (Safavian and Goring, 2013; Indriolo et al., 2014). TEM images revealed structures resembling autophagic bodies in the vacuoles of stigmatic papilla following self-incompatible pollinations (Safavian and Goring, 2013; Indriolo et al., 2014). In contrast, TEM images of compatible pollinations revealed vesicle-like structures in the stigmatic papillae under the compatible pollen contact site suggestive of secretory activity (Safavian and Goring, 2013; Indriolo et al., 2014). The presence of autophagosomes in *A. lyrata* self-incompatible pollinations was further confirmed using the GFP-ATG8a autophagosome marker (Safavian and Goring, 2013). Although these studies indicated that autophagy was activated following self-incompatible pollinations, it was unclear whether autophagy is essential for the rejection of self-incompatible pollen. Here, we have used two different self-incompatible (SI) transgenic *A. thaliana* lines (in the Col-0 and C24 accessions; Iwano et al., 2014; Zhang et al., 2019) to test the requirement of the autophagy system for self-pollen rejection. When loss-of-function mutants in the *AUTOPHAGY7* (*ATG7*) and *AUTOPHAGY5(ATG5)* genes (Chung et al., 2010; Ding et al., 2018) were combined with the SI-Col-0 and SI-C24 lines, we observed a partial breakdown of the self-incompatibility response in both accessions. This indicates that autophagic activity is required for the effective rejection of self-incompatible pollen, and thus, elucidates an additional downstream cellular event in the self-incompatibility pathway.

## Results

### Generation of self-incompatible *A. thaliana* transgenic lines carrying the *atg7* and *atg5* mutations to disrupt autophagy

To investigate the involvement of autophagy in self-incompatibility, two different transgenic *A. thaliana* SI lines were used to cross with *atg7* and *atg5* autophagy mutants. ATG7 and ATG5 are essential components in the formation of autophagosomes (reviewed in Ding et al., 2018), and both the *atg7* and *atg5* mutations lead to a loss of autophagy (Thompson et al., 2005; Chung et al., 2010; Young et al., 2019). The first self-incompatible line is in the Col-0 accession (SI-Col-0) and carries the *A. halleri SCR13, SRK13* and *ARC1* transgenes in a single T-DNA (*Ah-SCR13 Ah-SRK13 Ah-ARC1* #2 line; Zhang et al., 2019). This SI-Col-0 line was crossed to the previously characterized *atg7-2* mutant in the Col-0 accession (Chung et al., 2010). The F_1_ plants from this cross were selfed, and all pollination assays were carried out in the F_2_ generation. To confirm that any observed effects were not specific to the loss of *ATG7*, the previously characterized *atg5-1* mutation in the Col-0 accession (Thompson et al., 2005) was also crossed into the SI-Col-0 line, and taken to the F_2_ generation for pollination assays. The second self-incompatible line is in the C24 accession (SI-C24) and carries the *A. lyrata SCRb* and *SRKb* transgenes (SI-C24 #15-1 line; Iwano et al., 2015). As there were no *atg7* mutants available in the C24 accession, CRISPR/Cas9 technology was used to generate additional *atg7* mutants. Deletion of the *ATG7* gene was confirmed through PCR and sequencing of genomic regions flanking gRNA target sites (Supplemental Fig. S1). The *atg7-9* C24 mutant displayed the same overall wild-type plant and flower morphology (Supplemental Fig. S2) as seen for the *atg7-2* Col-0 mutant. The C24 *atg7-9* mutant was crossed with the SI-C24 line, and following selfing of the F_1_ plants, all pollination assays were carried out in F_2_ generation (T_4_ generation for the *atg7-9* mutation). For all three sets of F_2_ progeny, genotyping was first carried out to identify the transgenic plants carrying the SI transgenes, followed by genotyping for the *atg7* or atg5 mutations to identify plants that were wild-type or homozygous for the *atg* mutations.

### Disrupting autophagy breaks down the initial self-incompatible response of blocking pollen hydration

To assess the early stages of pollen rejection, the pollen hydration checkpoint was first investigated. Self-incompatible pollen from the parental SI-Col-0 or SI-C24 transgenic lines were used to pollinate pistils from the respective F_2_ progeny segregating for the *atg7* or *atg5* mutations, and pollen hydration assays were carried out by measuring the increase in pollen diameter at 10 min post-pollination (Lee et al., 2020). Positive controls of the SI-Col-0 and SI-C24 pollen placed on compatible wildtype Col-0 and C24 pistils, respectively, resulted in a large increase in diameter as a result of pollen hydration (Fig. 1A, B). In contrast, self-pollen placed on the SI-Col-0 and SI-C24 pistils resulted in very little pollen hydration at 10-minutes post-pollination (Fig. 1A, B). This lack of hydration was consistent between the SI-Col-0 and SI-C24 lines and indicative of the early recognition and rejection of self-incompatible pollen. Importantly, stigmas from the *atg7-2, atg5-1 and atg7-9* mutants showed normal SI-Col-0 and SI-C24 pollen hydration, respectively, indicating that disruption in the autophagic process did not perturb the compatible post-pollination responses (Fig. 1A, B).

**Figure 1.**
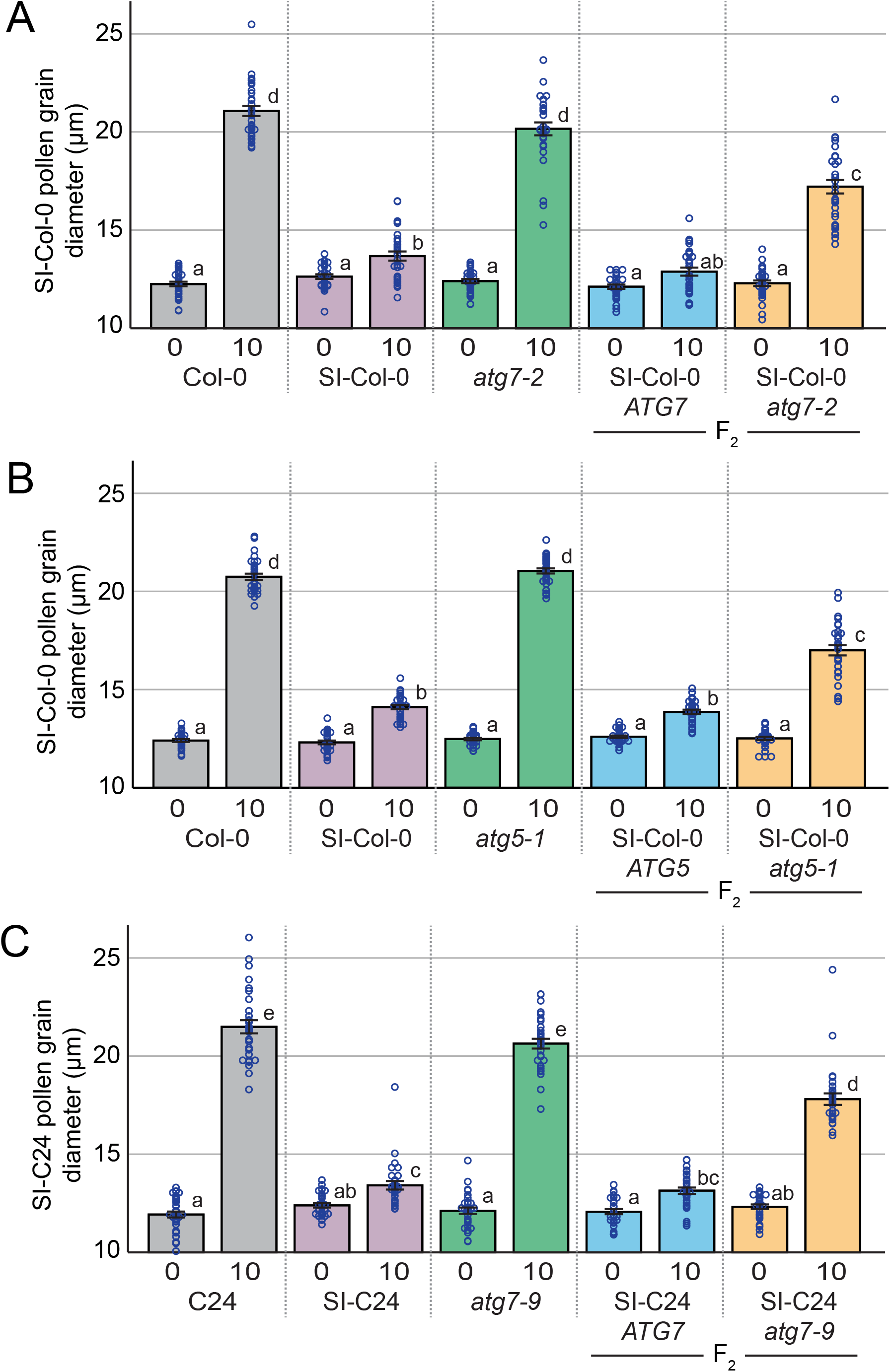
Pollen hydration assays on control and SI F2 progeny segregating for the *atg7* and *atg5* mutations. Pollen hydration assays for **A**, SI-Col-0 pollen grains placed on stigmas from Col-0, SI-Col-0, *atg7-2*, and F2 progeny from the SI-Col-0 x *atg7-2* cross; **B**, SI-Col-0 pollen grains placed on stigmas from Col-0, SI-Col-0, *atg5-1*, and F2 progeny from the SI-Col-0 x *atg5-1* cross; and **C**, SI-C24 pollen grains placed on stigmas from C24, SI-C24, *atg7-9*, and F2 progeny stigmas from the SI-C24 x *atg7-9* cross. Pollen hydration was measured by taking pollen grain diameters at 0- and 10-minutes post-pollination. Data is shown as Mean +/-SE with all data points displayed. n=30 pollen grains per line, P<0.05 (One-way ANOVA with Tukey-HSD post-hoc test).

For the F_2_ progeny from the SI-Col-0 x *atg7-2* cross, the SI-Col-0 plants homozygous for wildtype *ATG7* showed the same small increase in pollen hydration as the control self-incompatible pollinations at 10 min post-pollination. In contrast, when SI-Col-0 pollen was applied to the SI-Col-0 stigmas that were homozygous for the *atg7-2* mutation, an increase in pollen hydration was observed, indicating an incomplete rejection of the self-incompatible pollen (Fig. 1A). Similarly, pollen hydration assays on the F_2_ progeny from the SI-Col-0 x *atg5-1* cross showed that hydration of SI-Col-0 pollen on SI-Col-0 stigmas wild-type for *ATG5* displayed a small increase while SI-Col-0 pollen hydration on SI-Col-0 stigmas homozygous for the *atg5-1* mutation was significantly increased (Fig. 1B). A comparable set of pollen hydration experiments on the F_2_ progeny from the SI-C24 x *atg7-9* cross also showed a similar trend (Fig. 1C). SI-C24 stigmas that were wild-type for *ATG7* supported similar small increases in SI-C24 pollen hydration as the control self-incompatible pollinations, and the introduction of the *atg7-9* mutation into this background resulted in increased hydration of the self-incompatible pollen. Together, these results indicate that autophagy is required to fully prevent the hydration of self-incompatible pollen.

### Disruption of autophagy increases self-incompatible pollen tube penetration

With the observed increase in self-incompatible pollen hydration, the next stage of preventing self-incompatible pollen germination and pollen tube penetration was investigated. To determine if this block was affected by the *atg7* and *atg5* mutations, pollinated pistils were collected at 2- and 24-hours post-pollination and stained with aniline blue to visualize pollen tubes. At 2-hours post-pollination, SI-Col-0 and SI-C24 pollen was completely rejected by the respective self-incompatible pistils, and no pollen tubes were visible (Supplemental Fig. S2 and S3). Although we did not see any obvious difference at 2-hours post-pollination, we did observe an occasional pollen tube that penetrated the stigma barrier of the SI-Col-0 and SI-C24 *atg* homozygous mutants. To address whether we could observe a greater number of SI-Col-0 and SI-C24 pollen tubes after a longer period, pollinated pistil samples were then harvested at 24-hours post-pollination. A later time point following pollination is more reflective of the natural pollination process and provides more time for the self-incompatible pollen to germinate and grow into the pistil. Self-pollination of the SI-Col-0 and SI-C24 parental lines resulted in an absence of pollen tubes (Fig. 2A, 3A and 4A) while pollinations with compatible wild-type pollen led to an abundance of pollen tubes growing in the pistils (Fig. 2B, 3B and 4B). The results indicated that the SI-Col-0 and SI-C24 were able to fully reject self-incompatible pollen and accept wild-type compatible pollen at 24-hours post-pollination. The SI-Col-0 and SI-C24 pollen also produced numerous pollen tubes when placed on the *atg* mutant pistils, indicating that the absence of autophagy did not affect these compatible pollinations (Fig. 2C and 3C). The impact of the *atg* mutations on self-incompatibility at 24-hours post-pollination was examined by applying the SI-Col-0 and SI-C24 pollen on their respective F_2_ progeny homozygous for the *atg* mutations.

**Figure 2.**
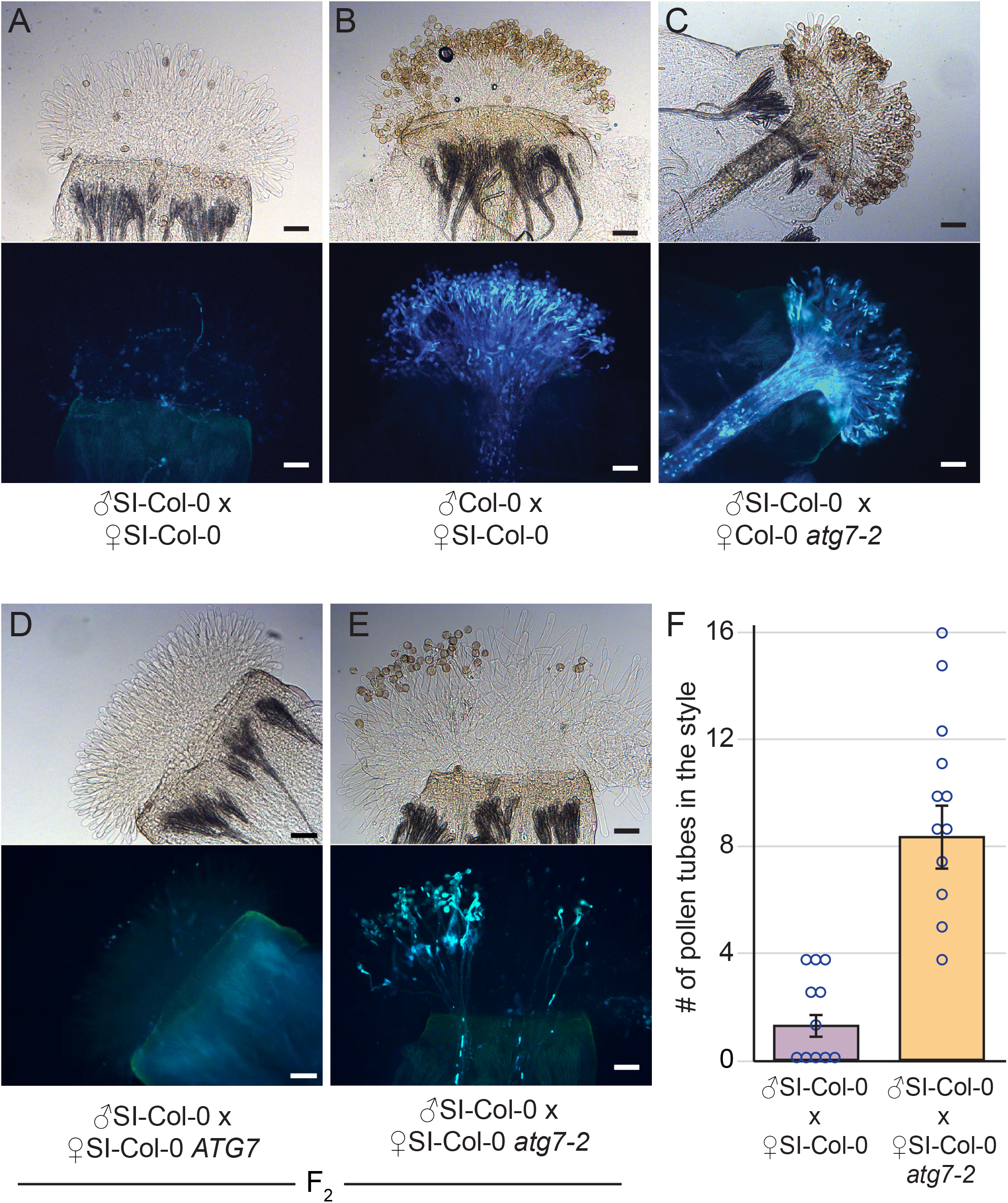
Representative images of aniline blue-stained SI-Col-0 or SI-Col-0 *atg7-2* pistils at 24 hrs following pollination. **A-E**, All stigmas were hand-pollinated with SI-Col-0 pollen or wild-type Col-0 pollen as indicated and collected at 24 hours post-pollination for aniline blue staining. The genotypes of the pistils are indicated for each panel. Pollen tubes penetrating through the stigma and style indicate a breakdown of self-incompatibility as seen in E. For each panel, top image = brightfield, bottom image = aniline blue. Scale bar = 100µm **F**, Bar graph showing the mean number of pollen tubes at 24 hrs post-pollination. Data is shown as Mean +/-SE with all data points displayed. n = 11-12 pistils per line. At 24 hours, pistils from SI-Col-0 *atg7-2* mutants allowed a significantly higher number of SI-Col-0 pollen tubes compared to the SI-Col-0 pistils (P<0.05).

**Figure 3.**
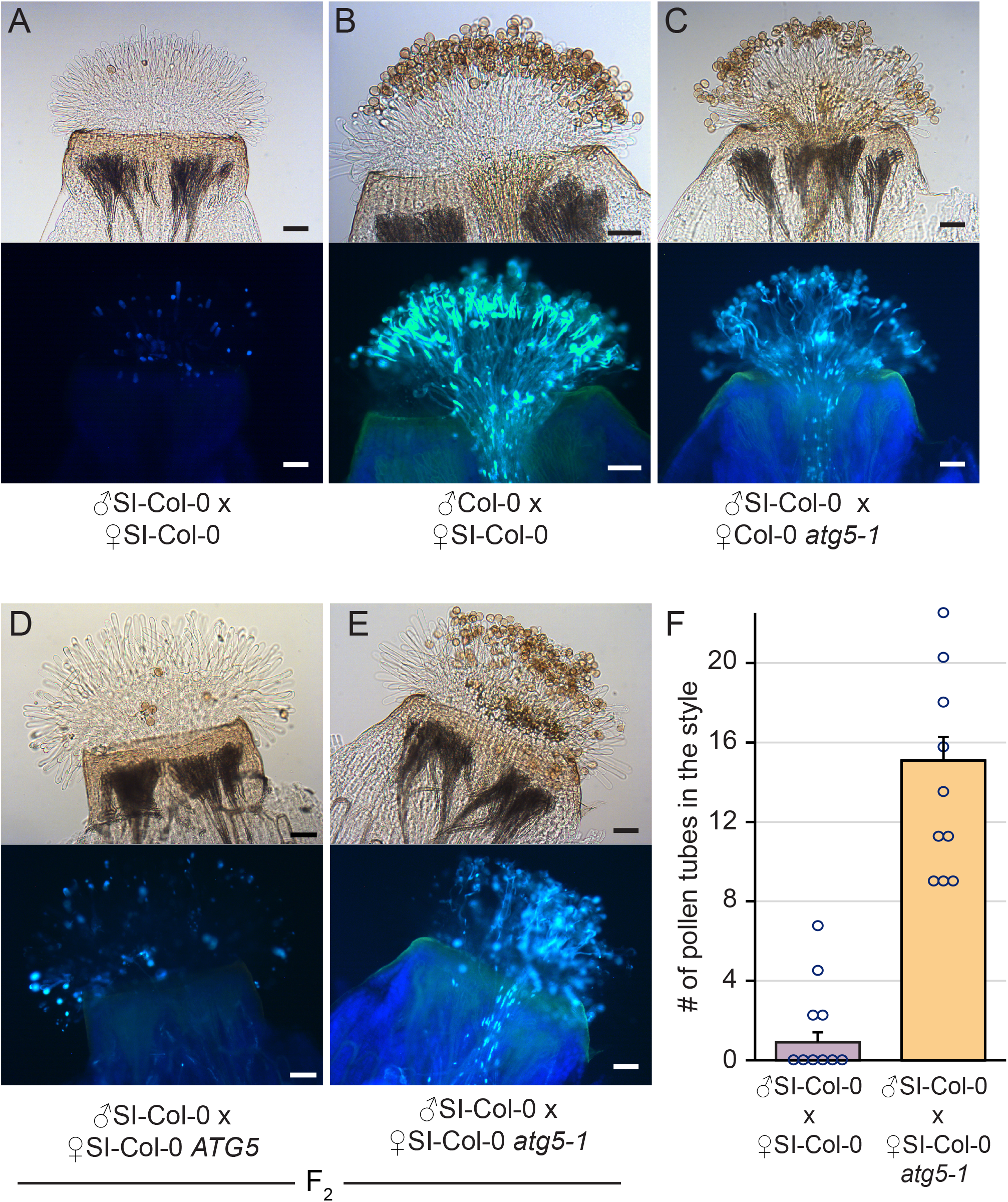
Representative images of aniline blue-stained SI-Col-0 or SI-Col-0 *atg5-1* pistils at 24 hrs following pollination. **A-E**, All stigmas were hand-pollinated with SI-Col-0 pollen or wild-type Col-0 pollen as indicated and collected at 24 hours post-pollination for aniline blue staining. The genotypes of the pistils are indicated for each panel. Pollen tubes penetrating through the stigma and style indicate a breakdown of self-incompatibility as seen in E. For each panel, top image = brightfield, bottom image = aniline blue. Scale bar = 100µm **F**, Bar graph showing the mean number of pollen tubes at 24 hrs post-pollination. Data is shown as Mean +/-SE with all data points displayed. n = 10 pistils per line. At 24 hours, pistils from SI-Col-0 *atg5-1* mutants allowed a significantly higher number of SI-Col-0 pollen tubes compared to the SI-Col-0 pistils (P<0.05).

When SI-Col-0 *atg7-2* stigmas or SI-Col-0 *atg5-1* stigmas were pollinated with SI-Col-0 pollen, there was a general increase in the number of pollen grains adhering to the stigma and forming pollen tubes when compared to the SI-Col-0 pistils (Fig. 2D-F and 3D-F). The pollinations were repeated for the SI-C24 F_2_ plants, and again the SI-C24 *atg7-9* pistils did show some SI-C24 pollen adhering to the stigmatic surface and successfully growing pollen tubes through the pistil while the SI-C24 *ATG7* pistils rejected the SI-C24 pollen (Fig. 4D-F). The number of pollen tubes were counted in the SI-Col-0 and SI-C24 pistils that were wild-type or homozygous for the *atg* mutations, and a significant increase in the number of pollen tubes were counted in the homozygous *atg7* and *atg5* mutants (Fig. 2F, 3F and 4F). Thus, the 24-hours post-pollination samples demonstrated that autophagy is required for fully blocking self-incompatible pollen germination and pollen tube growth through the self-incompatible pistil.

**Figure 4.**
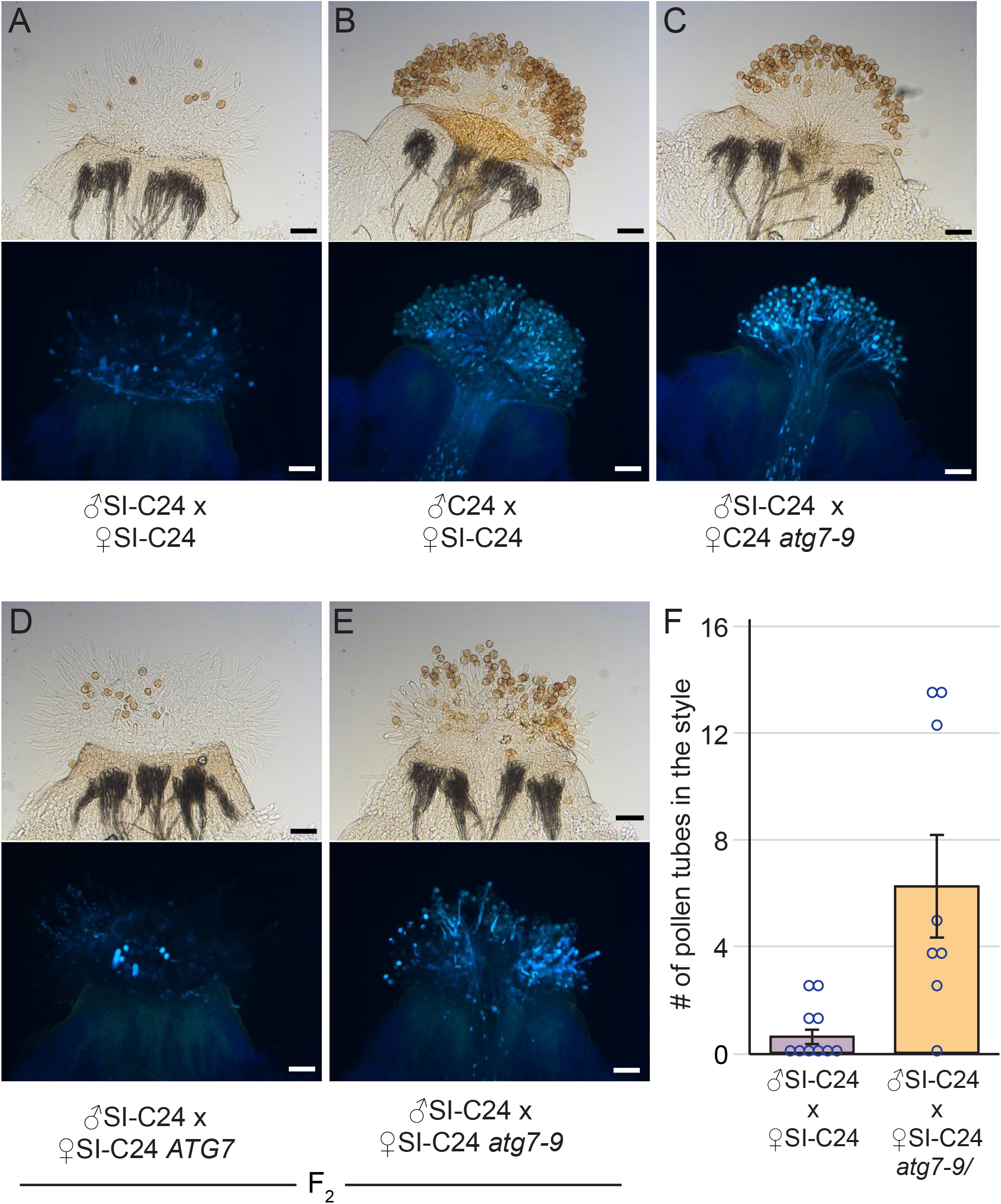
Representative images of aniline blue-stained SI-C24 or SI-C24 *atg7-9* pistils at 24 hrs following pollination. **A-E**, All stigmas were hand-pollinated with SI-C24 pollen or wild-type C24 pollen as indicated and collected at 24 hours post-pollination for aniline blue staining. The genotypes of the pistils are indicated for each panel. Pollen tubes penetrating through the stigma and style indicate a breakdown of self-incompatibility as seen in E. For each panel, top image = brightfield, bottom image = aniline blue. Scale bar = 100µm **F**, Bar graph showing the mean number of pollen tubes at 24 hrs post-pollination. Data is shown as Mean +/-SE with all data points displayed. n = 8-10 pistils per line. At 24 hours, pistils from SI-C24 *atg7-9* mutants allowed a significantly higher number of SI-C24 pollen tubes compared to the SI-C24 pistils (P<0.05).

### Loss of autophagy increases seed set in the self-incompatible lines

With the observation that self-incompatibility is partially lost in the SI-Col-0 *atg7-2*, SI-Col-0 *atg5-1* and SI-C24 *atg7-9* mutant pistils, resulting in more pollen tubes growing through the pistils, the impact of this on seed set was assessed. Manual pollinations were carried out using SI-Col-0 and SI-C24 pollen on pistils from the respective parental lines and F_2_ progeny homozygous for the *atg7* or *atg5* mutations, and 2-week old siliques were harvested for seed counting (Fig. 5; Supplemental Fig. S5). As expected for compatible pollinations in Col-0, C24, and the *atg* mutants, abundant seeds/silique were produced (Fig. 5A-C). As well, self-pollinations of SI-Col-0 and SI-C24 pistils resulted in a very limited seed yield consistent with the rejection of self-incompatible pollen (Fig. 5A-C). When the SI-Col-0 *atg7-2* pistils were pollinated with SI-Col-0 pollen, there was a significant increase in yield to 27 seeds/silique compared to 12.8 seeds/silique for SI-Col-0 indicating an incomplete rejection of the SI-Col-0 pollen by the SI-Col-0 *atg7-2* pistils (Fig. 5A). Similarly, when SI-Col-0 *atg5-1* pistils were pollinated with SI-Col-0 pollen, there was an increased average yield of 31.9 seeds/silique compared to 10.8 seeds/silique for SI-Col-0 (Fig. 4B). Finally, similar trends in seed yields were observed in the C24 accession. Self-incompatible pollinations with SI-C24 pistils resulted in a low average of 6.8 seeds/silique as expected yet when the SI-C24 *atg7-9* pistils were pollinated with SI-C24 pollen, there was a significant increase in seed yields to17.2 seeds/silique (Fig. 5C). Altogether, these results indicate that loss of autophagy was sufficient to disrupt the self-incompatibility pathway allowing for the growth of more SI pollen tubes through the SI-Col-0 and SI-C24 pistils carrying *atg7* or *atg5* mutations to initiate fertilization and seed production, and thus, confirming that autophagy is required for full rejection of self-pollen.

**Figure 5.**
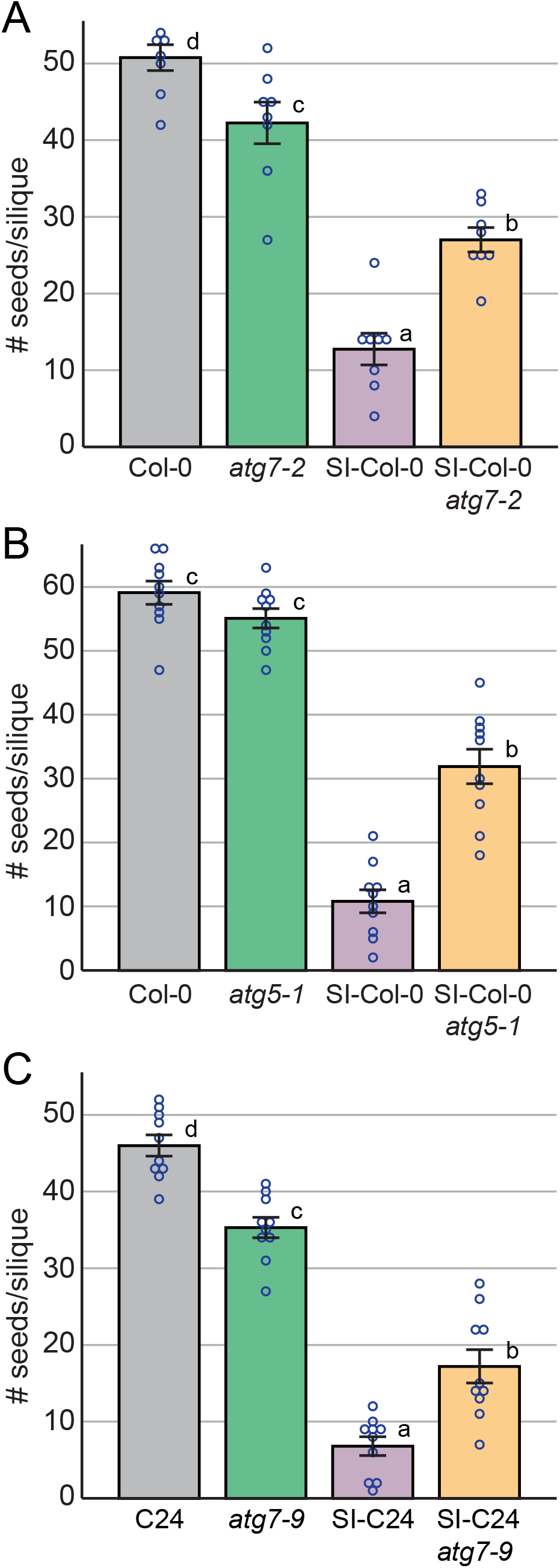
Seeds per silique counts for control and SI plants with the *atg7 or atg5* mutations. Seeds per silique counts following manual pollination with **A, B**, SI-Col-0 pollen or **C**, SI-C24 pollen on pistils with the indicated genotypes. Siliques were harvested at 2-weeks post-pollination and cleared for seeds counts. Data is shown as Mean +/-SE with all data points displayed. n= 8-10 siliques per line. Compared to the control SI pollinations, significant increases in the number of seeds per silique were observed for the SI plants that were homozyogous for the *atg7* or *atg5* mutations, P<0.05 (One-way ANOVA with Tukey-HSD post-hoc test).

### Autophagosomes are produced following self-pollination in the SI-C24 line

Previously, autophagosomes were detected by TEM following self-incompatible pollinations in *A. lyrata* and in the transgenic self-incompatible *A. thaliana* in the Col-0 accession transformed with *A. lyrata* S-locus genes (Safavian and Goring, 2013; Indriolo et al., 2014). To confirm that autophagosomes are also forming in the SI-Col-0 *Ah-SCR13 Ah-SRK13 Ah-ARC1* #2 line used in this study, *GFP-ATG8a* was used as an autophagosome marker (Thompson et al., 2005). Col-0 plants carrying the *ProUBQ10:GFP-ATG8a* construct (Kim et al., 2013) were crossed with the SI-Col-0 line, and the subsequent F_1_ generation was used for imaging. The SI-Col-0 *GFP-ATG8a* stigmas were imaged as unpollinated or manually pollinated with SI-Col-0 pollen. At 15-minutes post-pollination, many GFP punctae were observed in the SI-Col-0 *GFP-ATG8a* stigmatic papillae contacting SI-Col-0 pollen (Fig. 6A). Conversely, few puncta were observed in unpollinated SI-Col-0 GFP-ATG8a stigmatic papillae, consistent with a low background level of autophagy (Fig. 6C). The number of puncta were counted for both SI-Col-0 pollinated and unpollinated SI-Col-0 GFP-ATG8a stigmatic papillae (Fig. 6E), and there was a significant increase in the number of GFP-ATG8a-labelled punctae in the SI-pollinated SI-Col-0 *GFP-ATG8a* stigmatic papillae indicated a strong induction of autophagy. These observations confirm our previous findings that autophagy is activated in the stigmatic papillae with self-incompatible pollinations when compared to an unpollinated stigma (Safavian and Goring, 2013; Indriolo and Goring, 2014).

**Figure 6.**
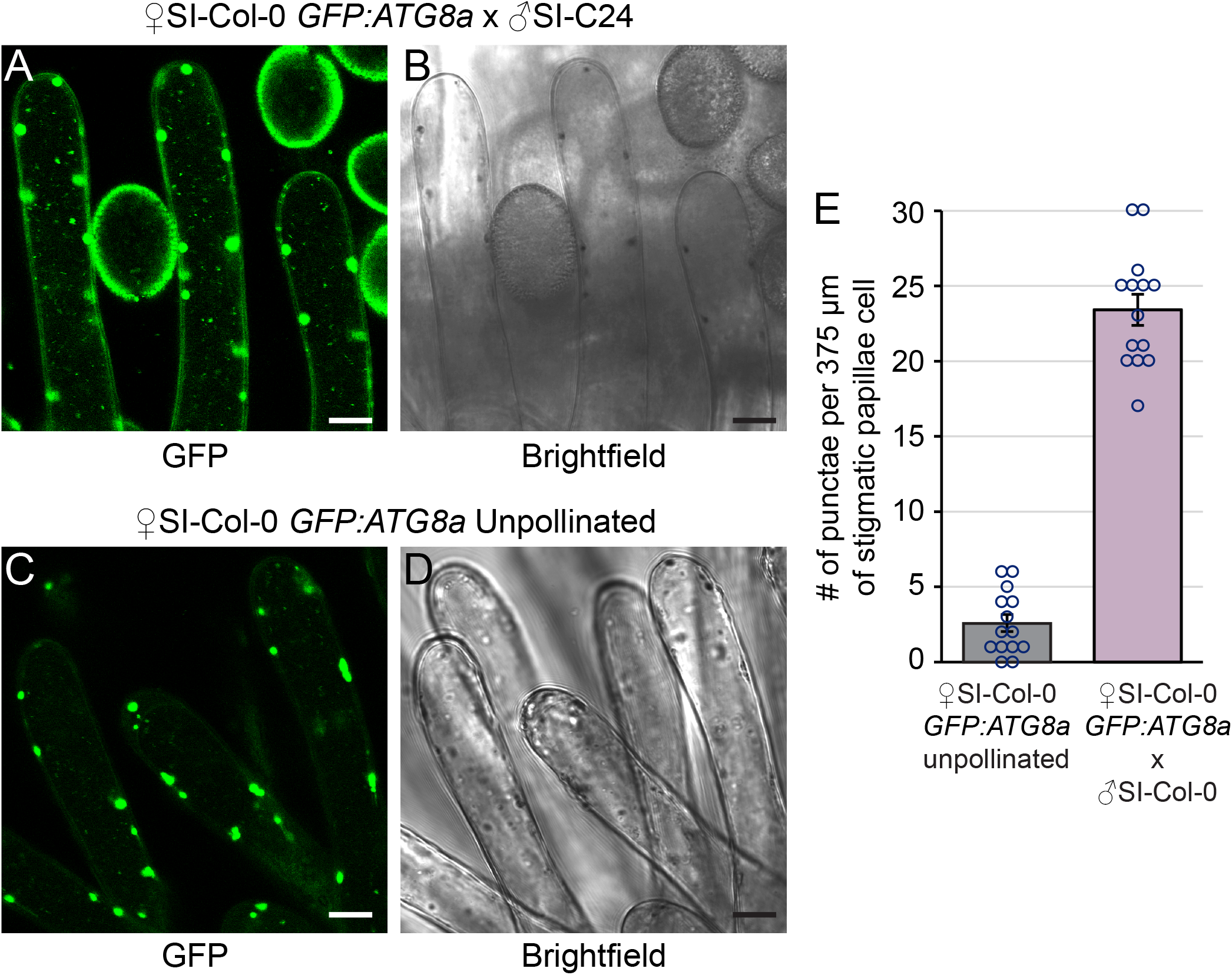
Autophagosomes are more abundant in SI-Col-0 *GFP-ATG8a* papillae following pollination with SI-Col-0 pollen. More autophagosomes marked by GFP-ATG8a are visible at 15-minutes post-pollination in the SI-Col-0 *GFP-ATG8a* papillae (**A-B**) compared to unpollinated SI-Col-0 *GFP-ATG8a* papillae (**C-D**). Scale bar = 10μm. **E**, Bar graph showing the mean number of autophagosomes present in each sample. Data is shown as Mean +/-SE with all data points displayed. n=14 papillae. At 15 min post-pollination with SI-Col-0 pollen, there is a significant increase in the number of GFP-labelled punctae in the SI-Col-0 GFP-ATG8a papillae (P<0.05). The large punctae are chloroplasts with autofluorescence and were not counted.

## Discussion

Self-incompatibility plays an important regulatory role in many outcrossing Brassicaceae species where producing inbred progeny would be disadvantageous (Shimizu and Tsuchimatsu, 2015; Durand et al., 2020). Previous studies have uncovered many important steps in the self-incompatibility pathway, but there are still outstanding questions on the cellular mechanisms by which self-pollen is rejected (reviewed in Doucet et al., 2016; Goring, 2017; Jany et al., 2019). In previous studies, we observed the presence of autophagosomes in the cytoplasm and autophagic bodies in the vacuole with self-incompatible pollinations in self-incompatible *A. lyrata* and transgenic self-incompatible *A. thaliana* Col-0 (Safavian and Goring, 2013; Indriolo and Goring, 2014). What was not known, however, was whether autophagy was necessary for self-incompatible pollen rejection. In this study, we addressed this question by crossing two different autophagy-deficient mutants (*atg7* and *atg5*) into two different transgenic self-incompatible *A. thaliana* lines, and demonstrated that a functioning autophagy system is required in the stigma for the full rejection of self-incompatible pollen. It is important to note that disrupting autophagy did not cause a full loss of self-incompatibility indicating that other cellular responses are required in conjunction with autophagy for self-incompatible pollen rejection. Importantly, this partial breakdown in the stigma was seen for all stages from increased pollen hydration to more pollen tubes growing through pistil to increase seed yields. Finally, the requirement of autophagy in the rejection of self-incompatible pollen was consistent between the three different autophagy deficient mutants *(<u>atg7-2, atg7-9</u>* and *atg5)* and the two different *Arabidopsis thaliana* accessions (Col-0 and C24) used in the SI lines indicating that there is a shared mechanism for self-incompatible pollen rejection.

The rationale for using both the SI-Col-0 and SI-C24 lines was to determine if there were any accession- or transgene-specific associations with autophagy. The SI-Col-0 carries the *A. halleri SCR13, SRK13* and *ARC1* transgenes (Zhang et al., 2019) while the SI-C24 line carries the *A. lyrata SCRb* and *SRKb* transgenes (Iwano et al., 2015). In addition to two different S-haplotypes from two different *Arabidopsis* species, the SI-Col-0 line also carries the *A. halleri ARC1* transgene. The fact that both accessions require an intact autophagy system in the stigma for full self-incompatible pollen rejection indicates that autophagy activation does not require ARC1, though the potential involvement of another related PUB E3 ligase cannot be ruled out (Indriolo et al., 2012; Indriolo and Goring, 2016). Our results indicate that autophagy is a clear commonality in the self-incompatibility response between these two accessions, although variation may exist in the execution of this cellular function. The question then arises to the purpose of autophagy activation in self-incompatibility.

Self-incompatible pollen carries the signals to be recognized as both self-incompatible pollen and compatible pollen, and it has been hypothesized that self-incompatibility blocks compatible pollen responses in the stigma (Dickinson, 1995). With the working model that secretory activity in the stigma is required for compatible pollen acceptance (reviewed in Dickinson, 1995; Doucet et al., 2016; Goring, 2017), several of the self-incompatibility cellular responses such as EX070A1 inhibition (Samuel et al., 2009; Sankaranarayanan et al., 2015), a strong and rapid calcium influx (Iwano et al., 2015), and the absence of actin focalization (Rozier et al., 2020) are consistent with blocking secretion (reviewed in Goring, 2017). Autophagy could also be an additional step to clear any secretory activity that was initially activated. Another possibility is that autophagy is resetting the cellular program to reject self-pollen. (Rodriguez et al., 2020) recently demonstrated a role for autophagy in switching *A. thaliana* cell fate determination following treatments with different signals. Autophagy was shown to be required for this switch and was proposed to “clean-up” the cell to allow for the new cellular program to be established. In the context of self-incompatibility, autophagy activation in the stigma could be linked to reprogramming the cells from the default compatible pollen responses to the self-pollen rejection pathway.

In conclusion, this study now firmly places autophagy as part of the cellular mechanisms leading to self-incompatible pollen rejection. Future studies will need to focus on how autophagy is activated in the self-incompatibility pathway and whether there are specific cytoplasmic components targeted for autophagic destruction. Understanding the mechanisms underlying self-incompatibility is important as seed yield of many important crop species is directly tied to their reproductive success, and the ability to modulate selfing would be a powerful tool in increasing yield.

## Materials and methods

### Plant materials and growth conditions

All *Arabidopsis thaliana* seeds (Col-0, C24, transgenic lines, T-DNA insertion mutants and CRISPR/Cas9 deletion mutants) were sterilized and cold stratified at 4°C for 3 days. Seedlings were germinated on ½ Murashige and Skoog (MS) medium plates supplemented with 0.4 (w/v) phytoagar (pH 5.8) and 1% (w/v) sucrose. Seedlings were grown at 22°C under 16h light for 7 days, and then transferred to Sunshine #1 soil supplemented with Plant Prod All Purpose Fertilizer 20-20-20 (1g/L water). All plants were grown in chambers under a 16-hour light/8-hour dark growth cycle at 22°C, with humidity kept under 50%.

For the Col-0 accession experiments, the *Ah-SCR13 Ah-SRK13 Ah-ARC1* #2 Col-0 line (Zhang et al., 2019) was crossed to the *atg7-2* mutant (Col-0 accession; Chung et al., 2010) and *atg5-1* mutant (Thompson et al., 2005). For the C24 accession experiments, the *SI-C24 #15-1* line (with *Al-SCRb AlSRKb*; Iwano et al., 2015) was crossed to the *atg7-9* mutant (C24 accession; see below). All transgenic lines were genotyped using primers found in Supplemental Table 1. For the C24 accession, plants were genotyped for *AlSCR*_*b*_, *AlSRK*_*b*_ and *atg7-9*. Progeny were analyzed in the F_2_ generation for all three crosses. For SI-Col-0 *GFP-ATG8a*, the *Ah-SCR13 Ah-SRK13 Ah-ARC1 #2* Col-0 line (Zhang et al., 2019), was crossed into the *ProUBQ10:GFP-ATG8a* line (Col-0 accession; Kim et al., 2013), and the resulting F_1_ progeny was used for all imaging.

### Plasmid construction and plant transformation

The *atg7-9* CRISPR/Cas9 deletion mutation was generated in C24 using previously described methods in the Col-0 accession (Doucet et al., 2019; Doucet et al., 2019). The final pBEE401E vector contained two guide RNAs targeting exons 3 and 8 of *ATG7* (Supplemental Fig. S1). *Arabidopsis thaliana* C24 plants were transformed using floral dip (Clough and Bent, 1998), and plants were dipped twice, 3 days apart, to maximize transformation efficiency. T1 seeds were collected and germinated on soil, and one-week old seedlings were sprayed with BASTA herbicide to select for transformants. PCR was used to confirm transformants (BASTA marker) and the presence of *atg7* mutations (deletions of the exon 3-8 region; Supplemental Figure S1). These *atg7* deletion mutants were then verified through sequencing of regions flanking the deletions. The *atg7* mutants showed a wild-type plant and flower morphology, along with the shorter life span typically seen in *atg* mutants (Thompson et al., 2005; Avila-Ospina et al., 2014). One homozygous mutant, *atg7-9*, from the T2 generation was crossed to the *SI-C24 #15-1* line, and all data was collected in the F_2_ generation of this cross (F_4_ generation for *atg7-9*).

### Confocal Microscopy

*A. thaliana* SI-Col-0 plants carrying the *ProUBQ10:GFP-ATG8a* construct were manually pollinated using pollen from the self-incompatible SI-Col-0 line. Pistils were removed at 10-minutes post-pollination and images of autophagosomes were taken at 15-minutes post-pollination. All images were taken using a Leica SP8 confocal microscope, with standard GFP excitation-emission wavelengths (488nm excitation, 495-545nm emission) with a pinhole diameter of 2.0AU. Leica LAS AF lite in combination with ImageJ was used for image processing and puncta counting. GFP punctae were counted in stigmatic papillae cell areas directly adjacent to SI-Col-0 pollen, and an equivalent cell area in unpollinated samples.

### Assays for Pollen hydration, Aniline blue staining, and Seed set

All post-pollination assays used manual pollinations and were conducted as described in Lee et al., 2020). For the SI-C24 plants, late stage 12 pistils were first emasculated and wrapped in plastic wrap overnight prior to pollination. For the SI-Col-0 plants, the pistils did not require emasculation due to the approach herkogamy phenotype (Zhang et al. 2019), and freshly opened flowers with unpollinated pistils were selected for these assays. All pollinations were completed at humidity levels under 50% for consistency. Pollen hydration assays were conducted by removing and mounting pistils on ½ MS, and lightly pollinating the stigmatic surface using a single anther. Images were taken at 0- and 10-min post-pollination using a Nikon sMz800 microscope, and pollen grains on the stigmatic surface were randomly selected for measurements using the NIS-elements imaging software. For the aniline blue staining, one anther was used to lightly pollinate each pistil, and the pistils were then collected at 2- and 24-hours post-pollination for staining as described in Lee et al., 2020). The aniline blue-stained pistils were mounted on a slide with sterile water, flattened, and imaged using a Zeiss Axioscope2 Plus fluorescent microscope. For seeds counts, pistils were manually pollinated with a single anther and were left for 10-14 days. The maturing siliques were then removed and cleared in 70% (v/v) ethanol for 3-5 days, or until seeds were visible (Beuder et al., 2020). Siliques were then mounted on a dry slide with the septum facing upwards and imaged using a Nikon sMz800 microscope, and seeds were counted using the NIS-elements imaging software.

## Supplemental Material

**Supplemental Figure S1**. Generation of the *ATG7* CRISPR/Cas9 deletion mutation in the C24 accession.

**Supplemental Figure S2**. Representative images of flowering plants for the SI-C24 and *atg7-9* mutant plants in the C24 accession.

**Supplemental Figure S3**. Representative images of aniline blue-stained pistils at 2 hrs following pollination with SI-Col-0 or Col-0 pollen.

**Supplemental Figure S4**. Representative images of aniline blue-stained pistils at 2 hrs following pollination with SI-C24 or C24 pollen.

**Supplemental Figure S5**. Representative images of cleared siliques used in seed counting for control and SI plants with the *atg7* mutation.

**Supplemental Table 1**. Primer Sequences.

## Acknowledgements

We thank Laura Canales and Arjun Sharma for technical assistance, and members of the Goring lab for critically reading this article. We are very grateful to Seiji Takayama (Nara Institute of Science and Technology; University of Tokyo) for providing the *SI-C24 #15-1* seeds (*A. thaliana* C24), Richard Vierstra (Washington University in St Louis) and Peter Bozhkov (Swedish University of Agricultural Sciences) for providing *atg7-2* and *atg5-1* mutant seeds (*A. thaliana* Col-0), and Marisa Otegui (University of Wisconsin-Madison) for providing the *ProUBQ10:GFP-ATG8a* seeds (*A. thaliana* Col-0).

