## Supplemental Figures for "Autophagy is required for the rejection of self-incompatible pollen in two accessions of transgenic *Arabidopsis thaliana*"

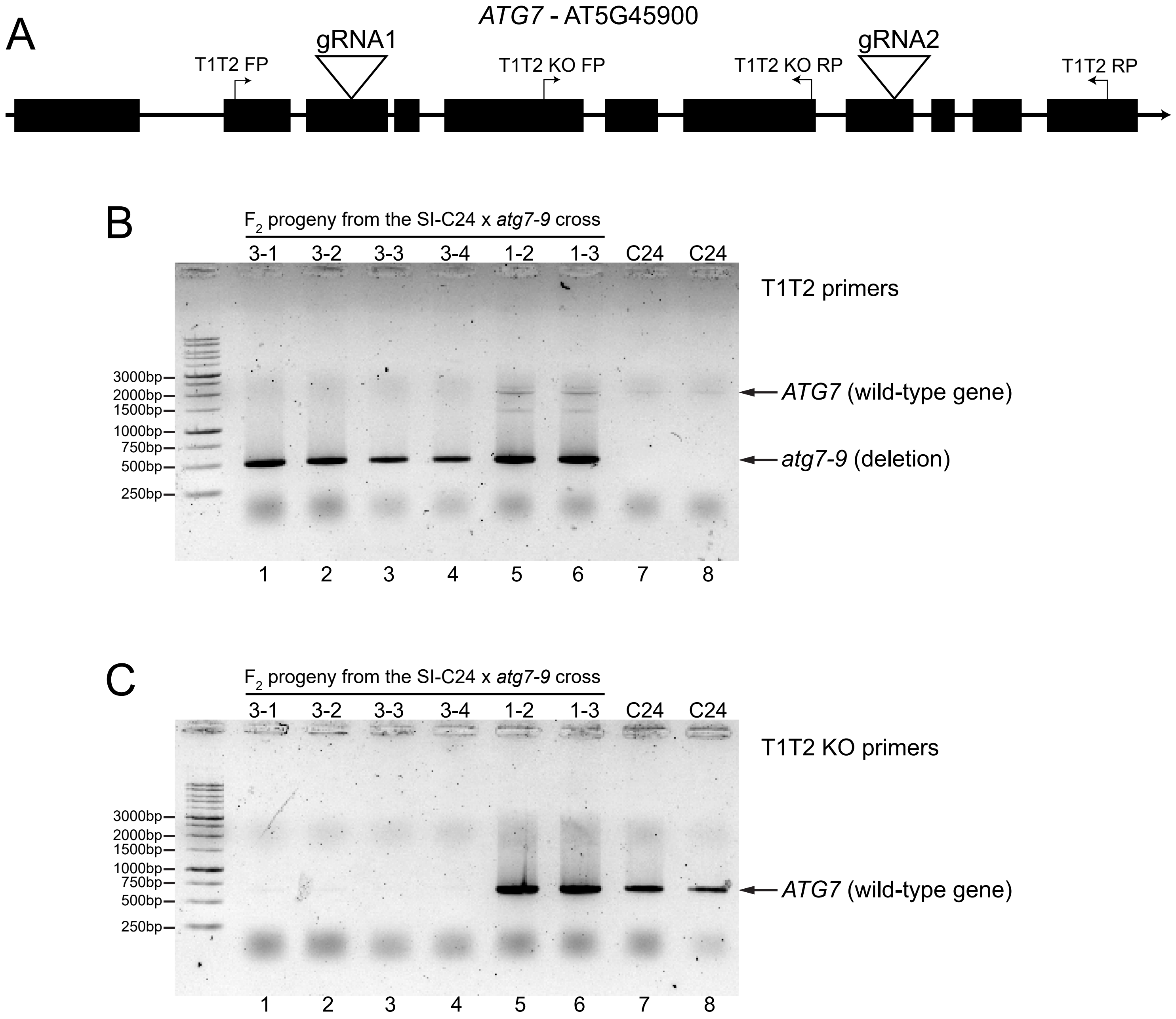


**Supplemental Figure S1. Generation of the *ATG7* CRISPR/Cas9 deletion mutation in the C24 accession.**

**A,** Schematic of the *ATG7* gene showing the location of the two gRNAs targeting exon 3 and exon 8 for the CRISPR/Cas9-mediated deletion in the C24 accession. The sites of the genotyping primers are also shown (T1T2 and T1T2 KO).

**B,** The *atg7-9* deletion was detected using primers flanking the gRNA cut sites (T1T2 FP, T1T2 RP). The top band is the wild-type *ATG7* gene and the bottom band is the *atg7-9* mutation (internal deletion of the region between exon 3 and exon 8).

**C,** Homozygotes for the *atg7-9* mutation were confirmed using primers in the internal region deleted in *atg7-9* (T1T2 KO FP, T1T2 KO RP). The $F_2$ progeny in lanes 1 to 4 are homozygous for the *atg7-9* mutation, while those in lanes 5 and 6 are heterozygotes. Lanes 7 and 8 are wild-type C24 plants.

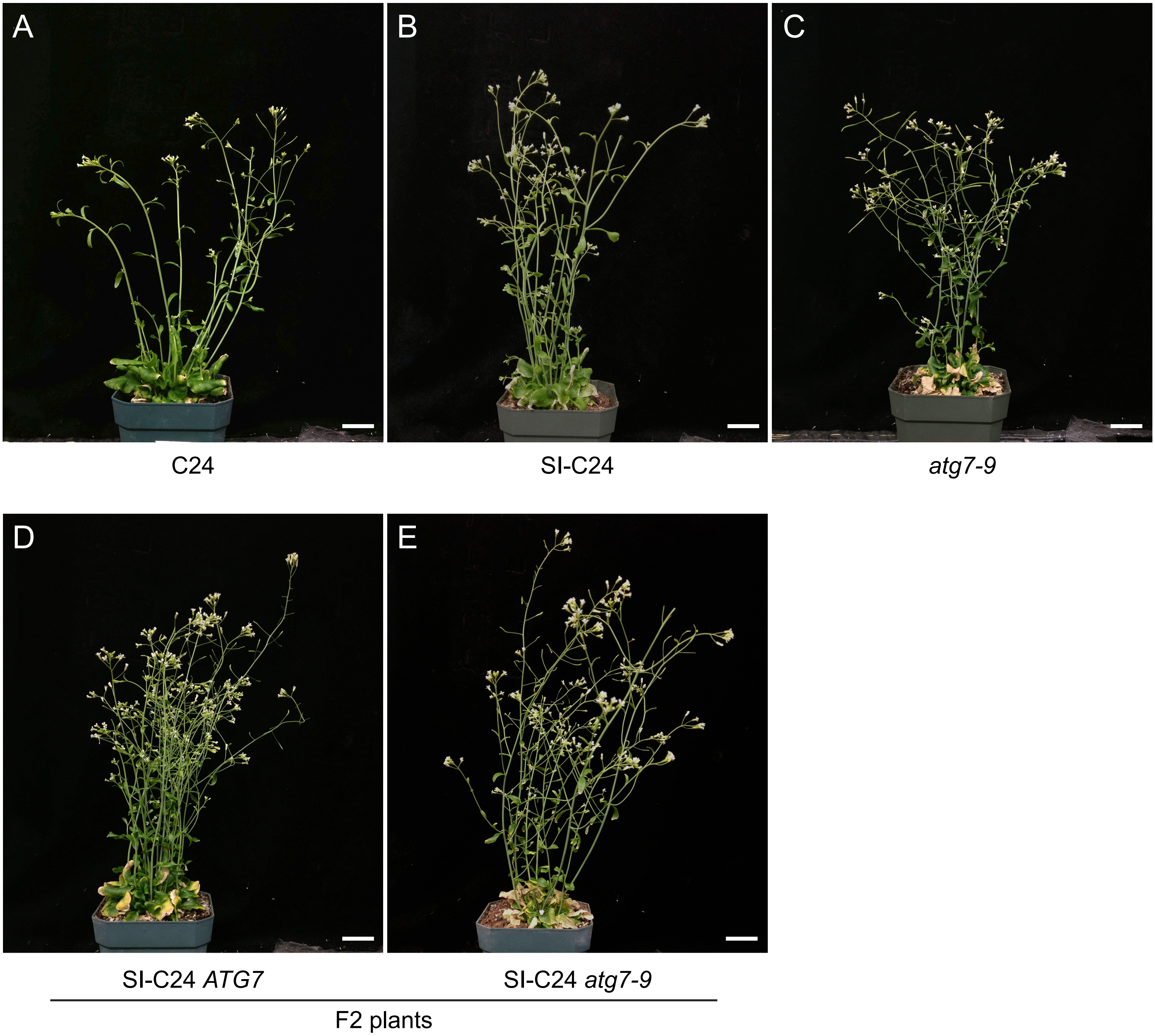


**Supplemental Figure S2. Representative images of flowering plants for the SI-C24 and *atg7-9* mutant plants in the C24 accession.**
**A-C,** shows wild-type C24, SI-C24 and *atg7-9* C24 mutant plants.
**D-E,** shows the $F_2$ plants from the SI-C24 x *atg7-9* cross. All plants shown are SI-C24 and segregating for the *atg7-9* mutation as indicated by the genotypes.
Scale bar = 2cm.

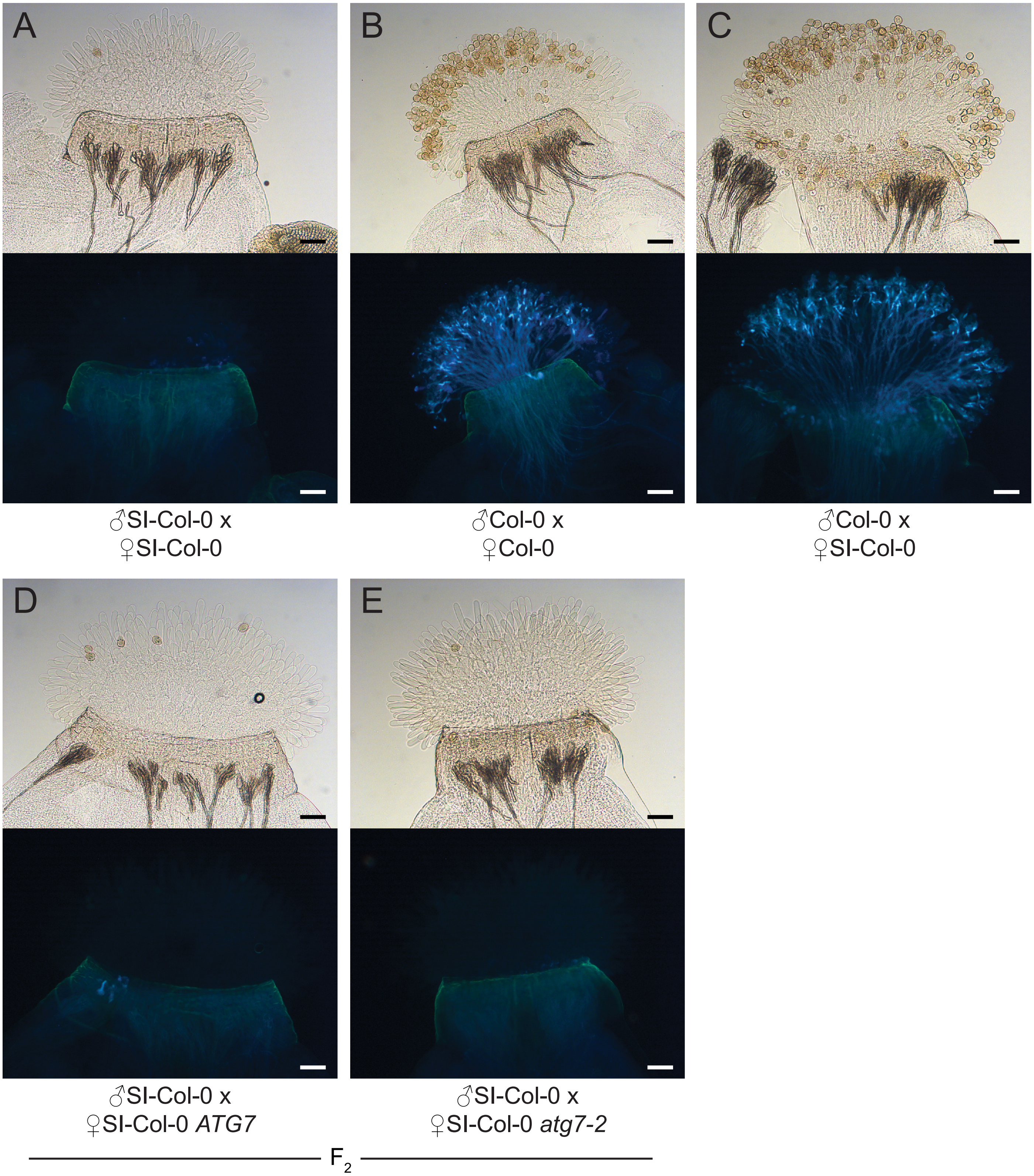


**Supplemental Figure S3. Representative images of aniline blue-stained pistils at 2 hrs following pollination with SI-Col-0 or Col-0 pollen.**

**A-F,** All stigmas were hand-pollinated with SI-Col-0 pollen or wild-type Col-0 pollen as indicated and collected at 2 hours post-pollination for aniline blue staining. The genotypes of the pistils are indicated for each panel.

For each panel, top image = brightfield, bottom image = aniline blue. Scale bar = 100μm.

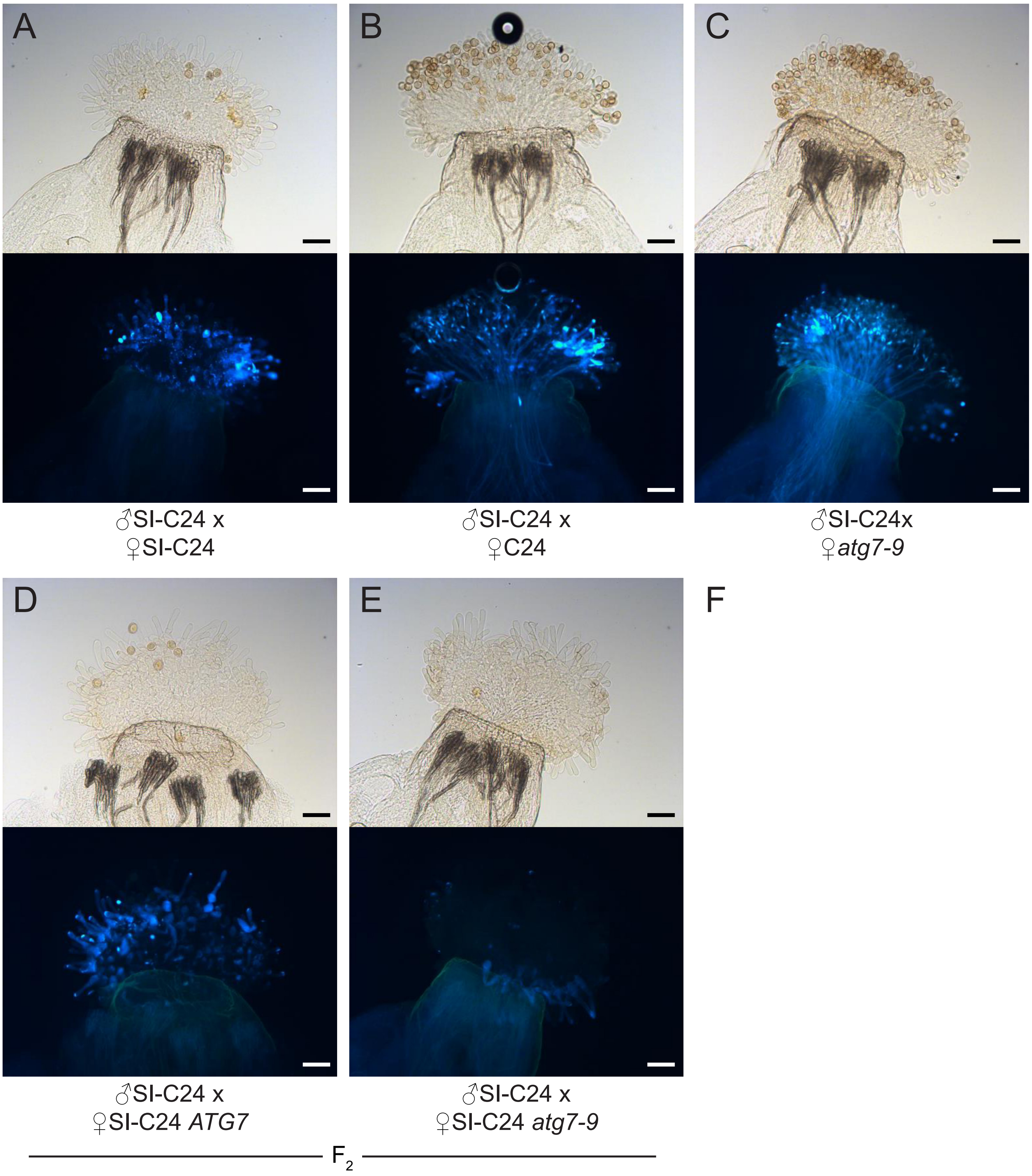


**Supplemental Figure S4. Representative images of aniline blue-stained pistils at 2 hrs following pollination with SI-C24 or C24 pollen.**

**A-F,** All stigmas were hand-pollinated with SI-C24 pollen or wild-type C24 pollen as indicated and collected at 2 hours post-pollination for aniline blue staining. The genotypes of the pistils are indicated for each panel.

For each panel, top image = brightfield, bottom image = aniline blue. Scale bar = 100μm.

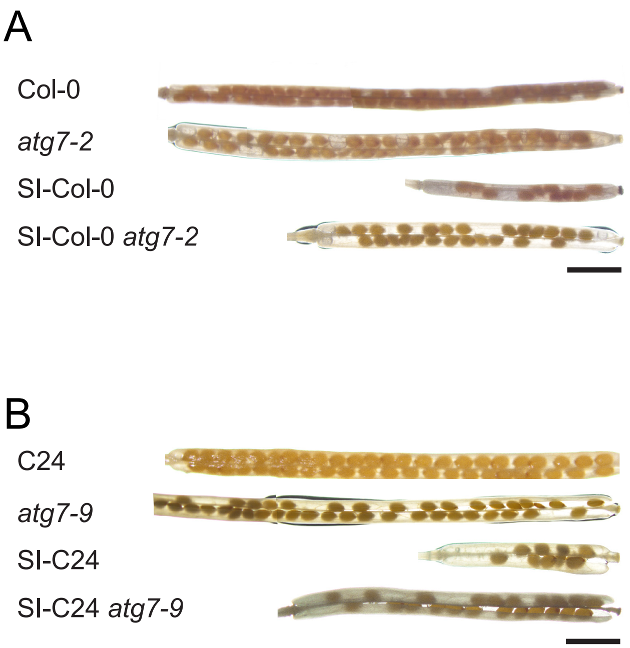


**Supplemental Figure S5. Representative images of cleared siliques used in seed counting for control and SI plants with the *atg7* mutation.**
Siliques were harvested at 2-weeks post-pollination following manual pollination with **A,** SI-Col-0 pollen or **B,** SI-C24 pollen on pistils with the indicated genotypes (Seed count graphs are shown in Figure 5).
Scale bar = 2 mm.

### Supplemental Table 1. Primer Sequences

| Primer Name | Primer Sequence | Used for |
|---|---|---|
| C24 atg7-9 target site 1 with BsaI and gRNA scaffold | ATTATTGGTCTCTAAACCTGCTTGTAATGATTGGCGCC AATCTCTTAGTCGACTCTACCAATA | Generating *atg7-9* deletion in C24 |
| C24 atg7-9 target site 2 with BsaI and gRNA scaffold | ATATATGGTCTCGATTGCATATCATCAGACAGAGAGG GTTTTAGAGCTAGAAATAGCAAGTTAAAAT | |
| C24 *atg7-9* T1T2 FP | CGTTCCCTGCGTTTGTACTT | Assessing *atg7-9* deletion |
| C24 *atg7-9* T1T2 RP | GAATTGTCGCCTTTTGCATT | |
| C24 *atg7-9* T1T2 KO FP | ATGAGATGGCGAGCATTACC | Assessing atg7-9 homozygosity |
| C24 *atg7-9* T1T2 KO RP | CATGGCGCATTACCATGTAG | |
| BastaR FP | CGGAGAGGAGACCAGTTGAG | Basta Resistance Screening |
| BastaR RP | GAGCTGGCAACTCAAAATCC | |
| *atg7-2* FP | TGCTAATTCCATGGATCCAACAAG | Detecting T-DNA insertion for *atg7-2* (Chung et al., 2010) |
| *atg7-2* RP | GAAGCCACCTAGTGAATATAGTCTATGGAC | |
| *atg7-2* RP-2 | TTGACCATCATACTCATTGCTGATCC | Assessing homozygosity of *atg7-2* (Chung et al., 2010) |
| ATG8-GFP-RT RP1 | GGTGAACTTCAAGATCCGCC | Genotyping GFP-ATG8a (Thompson et al., 2005) |
| GFP-ATG8a RP | ACTCATCCTTGCCTCGAGAGG | |
| Ah-ARC1 FP | CCGAAGTGCTACAGAGTGGA | Genotyping SI-Col-0 for SI components (Zhang et al., 2019) |
| Ah-ARC1 RP | TCCGAATCAGAGAACGTGAG | |
| SCRb FP | ATGAGGAATGCTACTTTCTTC | Genotyping SI-C24 for Al-SCRb (Iwano et al. 2015) |
| SCRb RP | TAGCAAAATCTACAGTCGCATA | |
| SRKb FP | ACCAAGATTCACGGTTCAGG | Genotyping SI-C24 for Al-SRKb (Iwano et al. 2015) |
| SRKb RP | ACGCTGTTCATGTGTCGAAG | |
| *atg5-1* FP | ATACACTTTAGAGGATATCC | Genotyping T-DNA insertion for *atg5-1* |
| *atg5-1* RP | CTGTCTTAACTTGTCCAGCCA | |
| LB1 | GCCTTTTCAGAAATGGATAAATAGCC | |
